# Dorsomedial striatal dopamine ramps down during interval timing

**DOI:** 10.64898/2026.08.01.742215

**Authors:** Hannah R. Stutt, Alexandra S. Bova, Matthew A. Weber, Madison M. McMurrin, Nandakumar S. Narayanan

**Affiliations:** Department of Neurology, University of Iowa

## Abstract

Dopamine is involved in disorders that degrade cognition such as Parkinson’s disease, ADHD, addiction, and schizophrenia; however, it is unclear how dopamine modulates brain circuits involved in cognitive processing. We investigated this problem by recording dopamine during interval timing, an elementary task that requires executive functions to estimate an interval of several seconds by making a motor response. We harnessed the fluorescent dopamine sensor dLight1.3b to record relative dopamine dynamics in the mouse dorsomedial striatum, which integrates information from cognitive cortical circuits and is required for interval timing. We found that: 1) dopamine activity ramped down during interval timing prior to increasing at reward delivery; 2) dopamine ramping dynamics predicted interval timing behavior; and 3) dopamine ramping dynamics were distinct between male and female mice and affected by amphetamine, a potent modulator of dopamine. These data provide insight into how dopamine modulates striatal circuits during interval timing and help better understand how dopamine dynamics contribute to cognitive dysfunction in dopamine-related brain diseases.

## INTRODUCTION

Dopamine is a neuromodulator involved in brain disorders such as Parkinson’s disease, ADHD, addiction, and schizophrenia^1,2^. These diseases can involve cognitive dysfunction that impairs executive functions such as working memory and attention, which can be modulated by dopamine^3–6^. However, executive dysfunction remains largely untreated in these and other disorders, in part because it is unclear how dopamine modulates circuits that affect executive function in the frontal cortex and striatum.

We investigated this problem using an interval timing task in mice, which requires subjects to estimate an interval of several seconds by making a motor response and requires working memory for temporal rules as well as attention to the passage of time^7^. Interval timing is highly translational to human diseases^8–10^ and translates to rodent models^11,12^. Interval timing requires the dorsomedial striatum in rodents^13–15^, which is functionally analogous to the human basal ganglia caudate nucleus that integrates cortical input^16,17^. We recorded relative dopamine dynamics during interval timing using the fluorescent dopamine indicator dLight1.3b^18^. Dorsomedial dopamine is modulated by task events such as cues, movements, or rewards^19,20^, and has been shown to ramp up in advance of timed rewards^21–24^, controlling temporal judgements. However, it is unknown how dopamine is modulated during temporal intervals.

To capture dopamine dynamics during timing, we recorded dopamine fiber photometry during a mouse-optimized interval timing task that requires mice to switch nosepokes after approximately 6 seconds to receive a food reward after 18 seconds^25,26^. We report three main results. First, we found that dopamine ramped down over the temporal interval in advance of prominent increases at reward delivery. Second, the slope of this downward ramp predicted when mice switched nosepokes and thus their internal estimates of time. Third, interval-dependent ramping was distinct in male vs. female mice and when dopamine was modulated by amphetamine. These data provide insight into cognitive and sex-specific mechanisms in dorsomedial striatal dopamine dynamics, which are highly relevant for brain diseases that disrupt cognition.

## METHODS

### Rodents

All experimental procedures were performed in accordance with the University of Iowa Institutional Animal Care and Use Committee (#3052039). Here, we include 21 C57BL/6J mice from Jackson Labs (Bar Harbor, ME) obtained at 12-14 weeks of age. All mice were housed in a 12-hour light-dark cycle with *ad lib* access to water. Mice were all trained on an operant based interval timing task using identical procedures and required to have >15 trials for dLight fiber photometry; a total of 12 mice (8 females and 4 males) were used as our baseline cohort to record dopamine dynamics during the interval timing task, and 5 mice (8 females and 4 males) for amphetamine sessions.

### Interval timing “switch” task

We trained mice on an interval timing task in which mice must guide their behavior based on their internal perception of how much time has passed, described in detail previously^11,15,26,27^. Mice were trained on this task in standard operant chambers inside sound attenuating cabinets (MedAssociates, St. Albans, VT). The chambers contained a food hopper with a response port on both sides, and one response port on the back wall of the chamber. Nosepoke responses were detected from infrared beams in each port. A trial was initiated by a nosepoke at the back response port, which triggered cue lights above the two front ports and an 3-kHz tone at 72 dB. Short- and long-interval trials had identical cues. All trials were reinforced with 20 mg sucrose pellets (BioServ, Flemington, NJ). Short-interval and long-interval trials occurred at 50% probability throughout each session. Short interval trials were reinforced for the first response after 6 seconds at the designated “short port” (counterbalanced left or right). Long-interval trials were reinforced for the first response after 18 seconds at the designated “long port”. Since long- and short-interval trials have identical cues, during long trials the mouse will start responding at the short port until approximately 6 seconds and then make the time-based decision to “switch” ports to obtain the sucrose reinforcer. During switch trials, or long interval trials, the time when animals depart the short response port is considered the “switch response time” and measures rodent’s internal estimate of time for when to switch to the long response port (Figure 1A). This task was designed to only analyze long interval switch trials, as responses on short trials rapidly decrease after 6 seconds and reward delivery, generating skewed time response histograms. Two experimental sessions of testing data per mouse were collected and analyzed for each treatment condition; sessions with <2 completed switch trials were excluded. Experimental sessions lasted 60 minutes.

**Figure 1:**
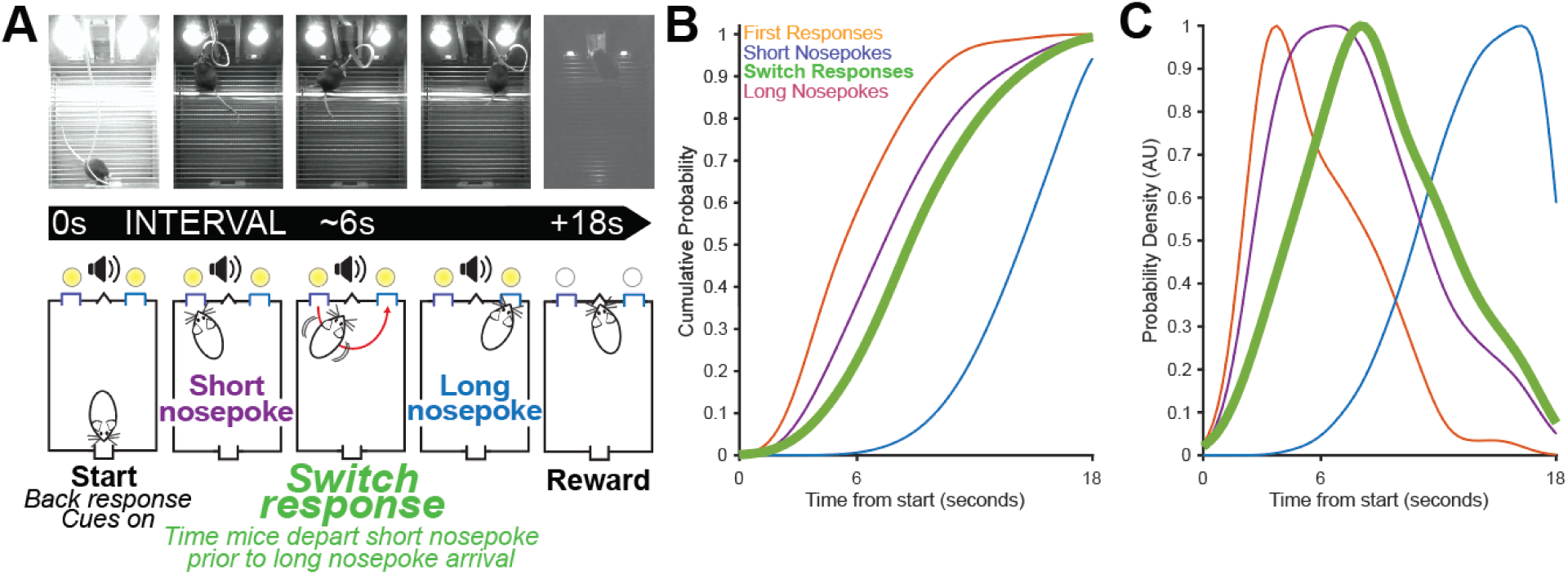
Mouse interval timing. (A) Mice initiate trials at a rear nosepoke, which triggers auditory and visual cues for the duration of the interval. Mice first respond at a short nosepoke (purple), and if no reward is delivered after approximately 6 seconds, they make a time-based decision to switch to the long nosepoke (blue) to wait for a reward after 18 seconds. The *switch response* (green) is defined as the moment mice depart the short nosepoke prior to arriving at the long nosepoke. (B) Cumulative probability of the behavioral metrics over the 18-second trial. The plot highlights the temporal distribution of first responses (orange), short nosepokes (purple), switch responses (green), and long nosepokes (blue). (C) Corresponding probability density (AU) of these behavioral responses plotted against the time from the start of the trial. Data from 12 mice (8 female).

### Drug administration

We observed how amphetamine affected interval timing behavior^28^ and dopamine dynamics in the dorsal striatum. *D*-amphetamine from Millipore-Sigma (A5880, Burlington, MA) was dissolved in sterile 0.9% saline. Sterile 0.9% saline was used as the vehicle solution and was injected intraperotineally 20 minutes before behavior. The next day, the same mice were injected with amphetamine at a dose of 1.5 mg/kg 20 minutes before behavior.

### Stereotaxic surgery

Mice were anesthetized with 4.0% isoflurane at 400 mL/min and maintained on 1.5%–3.0% isoflurane at 120 mL/min (SomnoSuite, Kent Scientific, Torrington, CT, USA). Craniotomies were drilled above the dorsomedial striatum (DMS; AP +0.5, ML +/- 1.4; counterbalanced left vs. right across mice). The virus dLight1.3b was infused for a total volume of 1 µL over 20 minutes (0.5 µL at DV -2.6 and DV -2.8). An optic fiber (4mm, Doric Lenses, Quebec, QC, Canada) was then implanted (DV -2.7; Figure 6A) in the DMS. At least three more craniotomies were drilled to insert skull screws (to anchor the headcap assemblies), which were sealed with cyanoacrylate (“SloZap,” Pacer Technologies, Rancho Cucamonga, CA) and accelerated with “ZipKicker” (Pacer Technologies) and methyl methacrylate (AM Systems, Port Angeles, WA). Following surgery, mice recovered for 1 week before food restriction and interval timing training.

### Fiber photometry recordings and analysis

After training in the interval timing task, mice were acclimated to the fiber photometry equipment and chambers. Mice were tethered to a unilateral optical patch cord connected to a Fiber Photometry System from Doric Lenses Inc (Quebec, Canada) with two excitation wavelengths (405 nm LED isosbestic signal modulation and 470 nm LED dopamine dependent dLight signal)^29^. LED light was emitted, and isosbestic and dLight fluorescence wavelengths were measured through integrated LED mini cubes and photodetector. Changes in dLight signal were aligned to task events (using Open Ephys). The raw dLight and isosbestic data was then down sampled to 100 Hz and passed through a lowpass filter (5 Hz) to de-noise recordings. To correct the recordings for photobleaching, we fit a double exponential decay function due to the initial decay in fluorescence and the more gradual decay in fluorescence through the recording. To correct for motion related fluorescence and autofluorescence, we correlated the dLight signal to the isosbestic signal and derived an interpolation line for each session. The interpolation line provided a slope and intercept for each isosbestic value; the estimated motion within each dLight value was then calculated with these inputs. The dLight signal was then normalized by z-scoring each session to compare activity across sessions and mice.

### Histology

Mice were sacrificed after completing fiber photometry experiments. Mice were anesthetized with ketamine (100 mg/kg ip) and xylazine (10 mg/kg ip) prior to transcardial perfusion. Perfusions were performed with phosphate buffered saline and 4% paraformaldehyde (PFA). Whole heads were removed and placed in fixed in 4% PFA for 24 hours. Brains were then extracted and placed back into 4% PFA for another 24 hours. To prepare the brains for cryosectioning, brains were transferred to a 30% sucrose solution for 48 hours and then frozen at -80 ℃ in cryoprotectant solution. Brains were sectioned using a cryostat (Lecia Biosystems, Deer Park, IL) and coronal or horizontal dorsal striatal sections at 40 µm were collected. Dorsomedial striatal sections with optic fiber tracts were selected and stained to visualize dLight and tyrosine hydroxylase expression. To stain for dLight expression, we used an anti-GFP primary made in rabbit and a goat anti-rabbit Alexa Fluor 488 secondary antibody. To stain for tyrosine hydroxylase, we used an anti-tyrosine hydroxylase primary made in chicken and a goat anti- chicken Alexa Fluor 568 secondary antibody. Stained sections were then mounted on slides and imaged using an Olympus VS120 slide scanning microscope (Olympus, Center Valley, PA; Figure 6A).

## Results

We trained 18 mice (10 male / 8 female) to perform an interval timing task identical to prior procedures^15,27,30^. Mice switched nosepokes at 8.9 +/- 0.2 seconds, with a coefficient of variation of 0.39 +/- 0.09 (CV; Fig 1B-C).

In these mice, we implanted fiber optics in the dorsomedial striatum along with dLight1.3b and recorded dopamine dynamics during interval timing (Fig 2A-B). In line with prior work, we found prominent modulations in dopamine dynamics as measured by z-scored dLight signals during trial start and reward in nearly every animal (Fig 2C-E)^19–21,31^. However, dLight activity around switch times appeared more heterogenous (Fig 2D).

**Figure 2:**
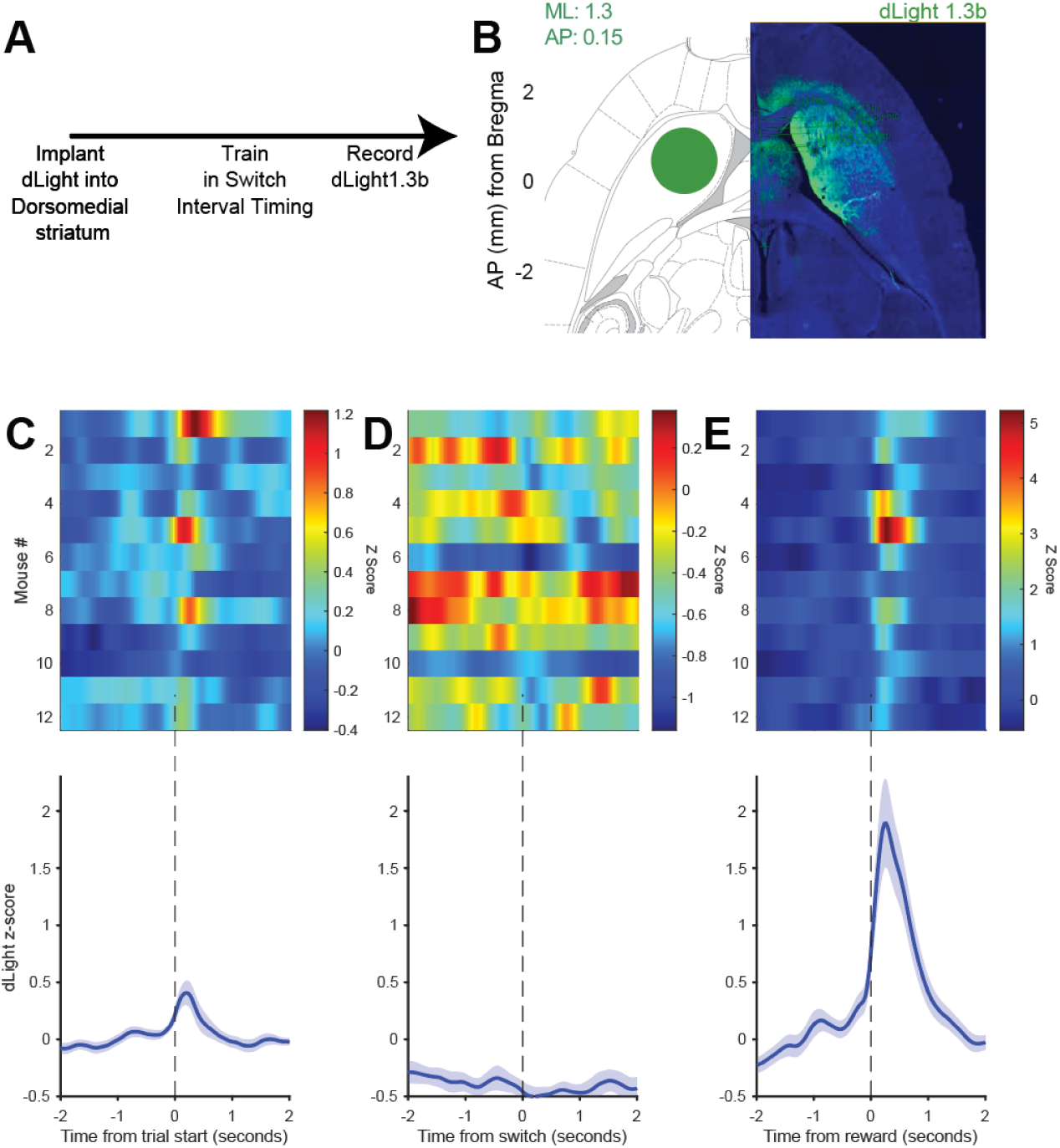
dLight activity during interval timing. (A) Experimental timeline detailing viral expression of the dopamine sensor, dLight1.3b, in the dorsomedial striatum, followed by behavioral training in the switch interval timing task, and fiber photometry recording. (B) Schematic representation of the target surgical coordinates within the dorsomedial striatum (AP: 0.15, ML: 1.3, DV: -2.6). (Right) Representative horizontal histological section confirming dLight1.3b expression and fiber placement in the target region. (C–E) Heatmaps of dLight z- score dynamics from 18 individual mice and the corresponding population average traces. For the traces. Solid lines represent the group mean, and shaded regions indicate ± SEM. (C) is aligned to trial start, (D) is aligned to the switch response, and (E) is aligned to reward.

We analyzed dLight dynamics over the interval. Over the 18-second interval timing trial, dLight signals decreased in nearly animal (Fig 3A-B). On average, the minimum time of the dLight signal was 9.83 seconds (Fig 3B). To further quantify these dynamics, we analyzed data from every single trial from every mouse. We found that there was a main effect of time in the interval on dLight signal (F_(1,5305)_ = 17.6, *p* =0.00002), a main effect of switch time (F_(1,5305)_ = 4.8, *p* =0.03), and a trend towards an interaction (F_(1,5305)_ = 3.6, *p* =0.06). Taken together, these data suggest that dLight activity was affected by time in the interval as well as switch time.

**Figure 3:**
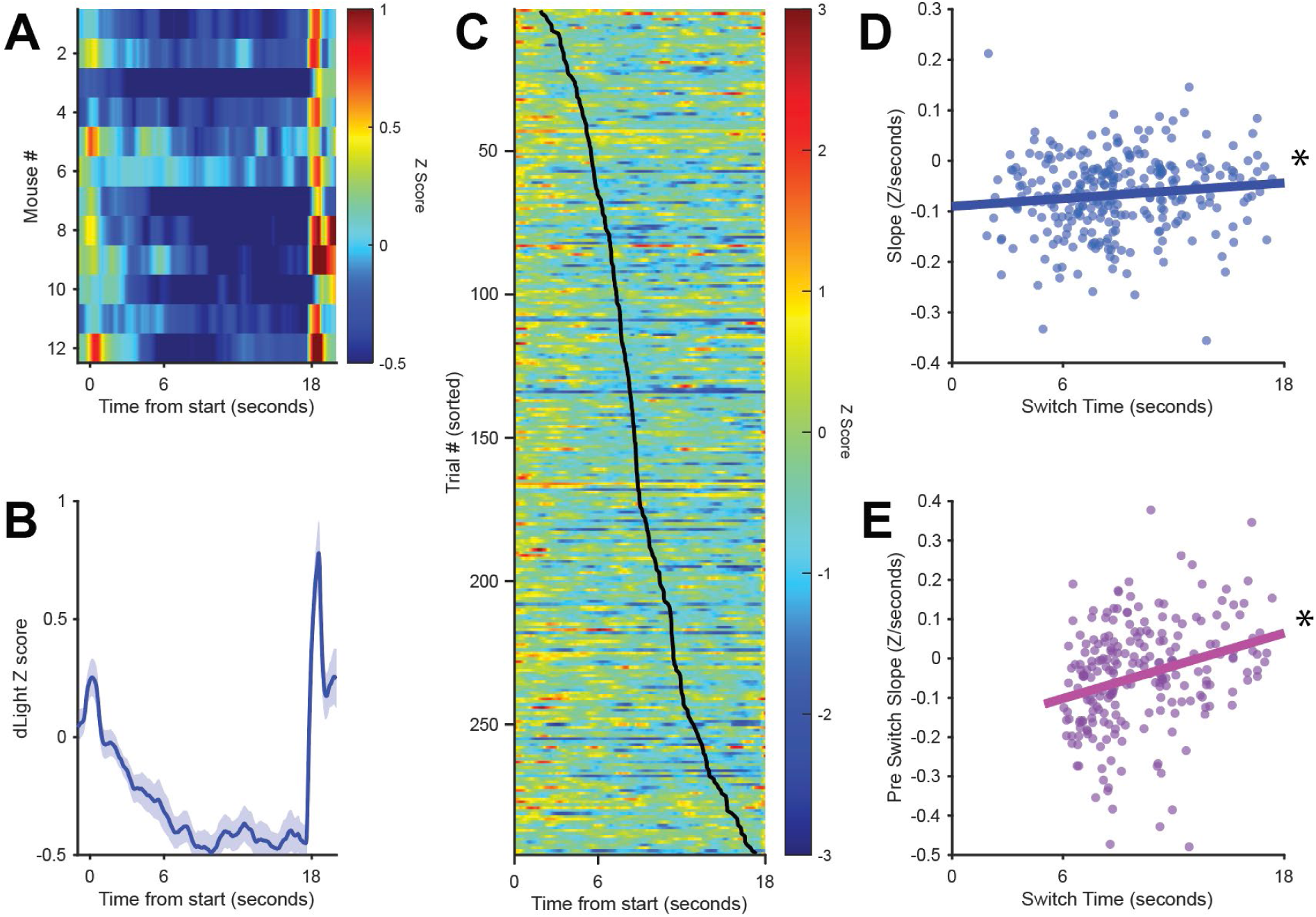
Dopamine (dLight) dynamics and correlations with behavioral switch times. (A) Heatmap illustrating the per-animal average dLight z-score for 11 individual mice over an 18- second trial period. The color scale denotes z-scores ranging from -0.5 (dark blue) to 1.0 (dark red). (B) Population average trace of the dLight z-score relative to the time from the start of the trial. Solid lines represent the group mean, and shaded regions indicate ± SEM. (C) Heatmap displaying dLight z-scores for all single trials across all animals, sorted by behavioral switch time (overlaid black ticks). (D) Scatter plot demonstrating the relationship between the initial dLight signal slope (measured from 0 to 10 seconds) and the switch time. (E) Scatter plot showing the relationship between the dLight signal slope in the period just before the behavior (measured from -4 seconds to the switch) and the switch time. * = p < 0.05 via linear mixed-effects models. Data from 12 mice.

To explore this in depth, we examined changes in dLight activity at a trial-by-trial level (Fig 3C). We noticed that on trials with an earlier switch time, dLight signals were flatter relative to trials with a later switch time, on which dLight signals seemed to ramp down^21,23^. We calculated the change in dLight activity from trial start to the average minimum over trials 9.8 seconds. A linear mixed-effects model accounting for mouse-specific effects revealed a main effect of dLight slope on switch time (F_(1,292)_ = 4.6, *p* = 0.02; Fig 3D).

To make sure that this was not related to trial averaging or other phenomena, we aligned dLight activity around switch time and examined dLight slope 6 seconds prior to switch responses. We found that dLight slope prior to switch time was strongly predictive of switch time (F_(1,292)_ = 67.1, p = 10^-15^). This held when switch responses <6 were excluded (F_(1,227)_ = 21.4, p =0.000006). These data suggest that dLight dynamics were related to switch time, which reflect animals’ internal estimates of time. Taken together, these data provide strong evidence that dopamine ramps down over the interval and is predictive of interval timing behavior.

Striatal dopamine dynamics are different in males and females^32–35^. We examined if these differences affected dLight dynamics in male vs. female mice during interval timing. As with prior work, there was no difference in interval timing behavior between male and female mice (8 female, 5 male; mean switch time; *p* = 0.37, CV *p* =0.57; Fig 4A-B). dLight dynamics are shown in Figure 4C-E. For the epoch prior to 9.83 seconds, a trial-by-trial linear mixed-effects model of dLight activity over the interval revealed a main effect of time (F_(1,2651)_ = 152.4, *p* < 10^-16^) and a main effect of sex (F_(1, 2651)_ = 3.9, *p* =0.04), and an interaction between time and sex (F_(1, 2651)_ = 5.8, *p* = 0.02; Fig 4E). These data provide evidence that males and females may rely on different dorsal striatal dopamine dynamics during interval timing.

**Figure 4:**
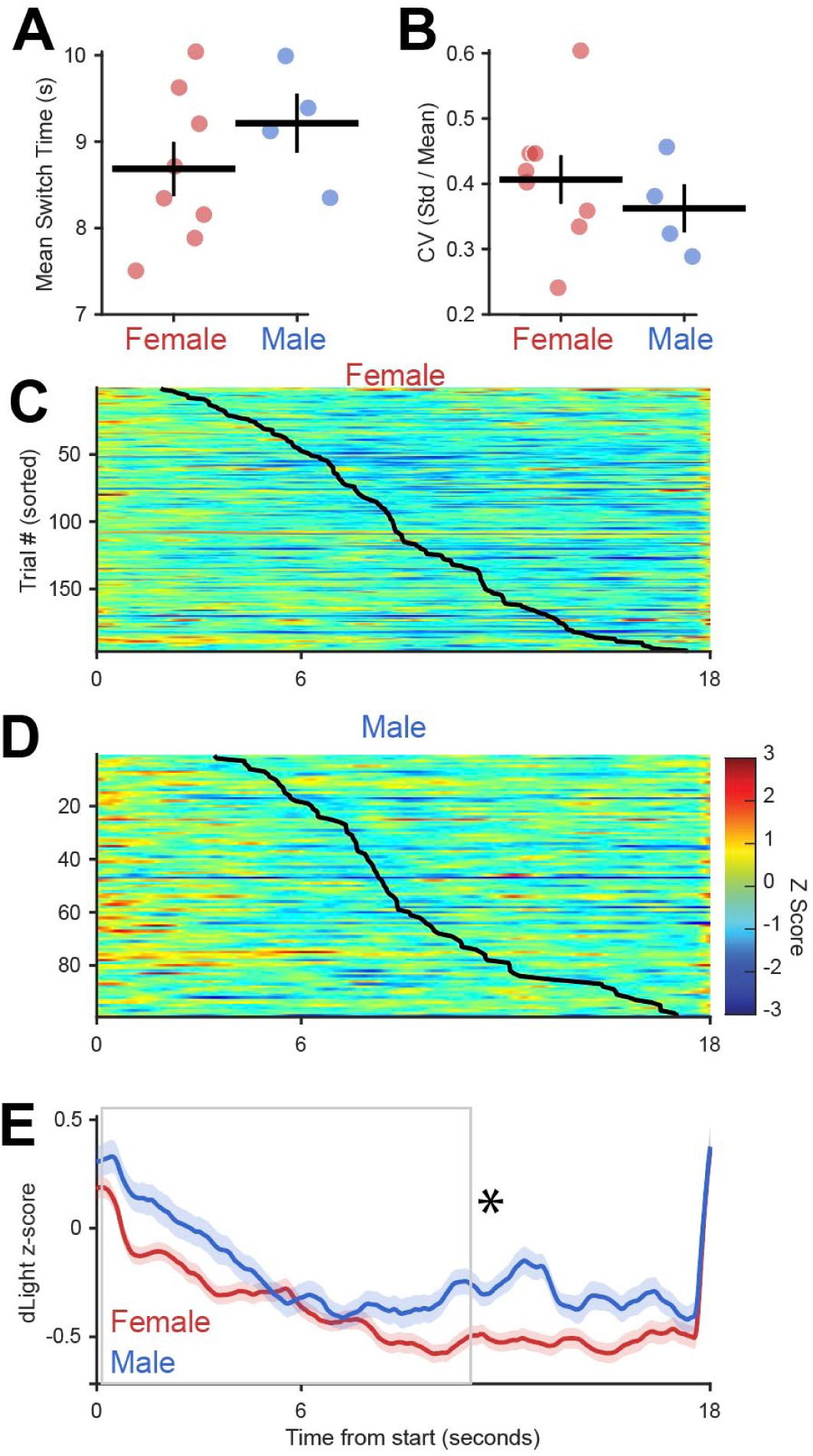
Interval timing dynamics in male vs. female mice. (A) Mean switch time (seconds) and (B) switch time coefficient of variation (CV) for female (magenta) and male (light blue) mice. Solid black horizontal and vertical lines indicate the group mean and standard error of the mean (SEM), respectively. (C) Heatmap of normalized dLight fluorescence (z-score) across all single trials from female mice and (D) from male mice. The x-axis represents time from the start of the trial (0–18 seconds). Trials on the y-axis are sorted sequentially by switch time, which is denoted by the overlaid solid black line. (E) Average dLight signal (z-score) across the 18- second trial window for female (magenta) and male (light blue) mice. Solid lines represent the group mean, and shaded regions indicate ± SEM. * = p < 0.05 via linear mixed-effects models. Data from 12 mice.

We analyzed how perturbing dopamine with amphetamine affected dopamine dynamics. Amphetamine powerfully reverses dopamine transport and re-uptake^28^. Notably, amphetamine can have complex effects on interval timing, affecting both temporal precision and accuracy, although there is considerable heterogeneity among studies, and there are small effects in some studies. In 5 animals that received saline and then amphetamine we found no differences in mean switch time (Fig 5A-B; 9.2 ± 0.7 vs 9.8 ± 0.7; *p* = 0.19) or CV (Fig 5C; 0.42 ± 0.04 vs 0.37 ± 0.05; *p* = 1). Furthermore, we found no differences in the timing of first nosepoke (Fig 5E; *p* = 0.44) or nosepoke duration (Fig 4E; *p* = 0.44, or traversal times between short and long nosepoke; Fig 4F; *p* = 0.44). While past studies have shown that amphetamine decreases temporal precision, data in this manuscript are derived from sessions with sufficient trials for dLight analysis (15+), and null effects have been reported in past work. The equivalent effects here enable us to compare dopamine dynamics when behavior is matched between sessions.

**Figure 5:**
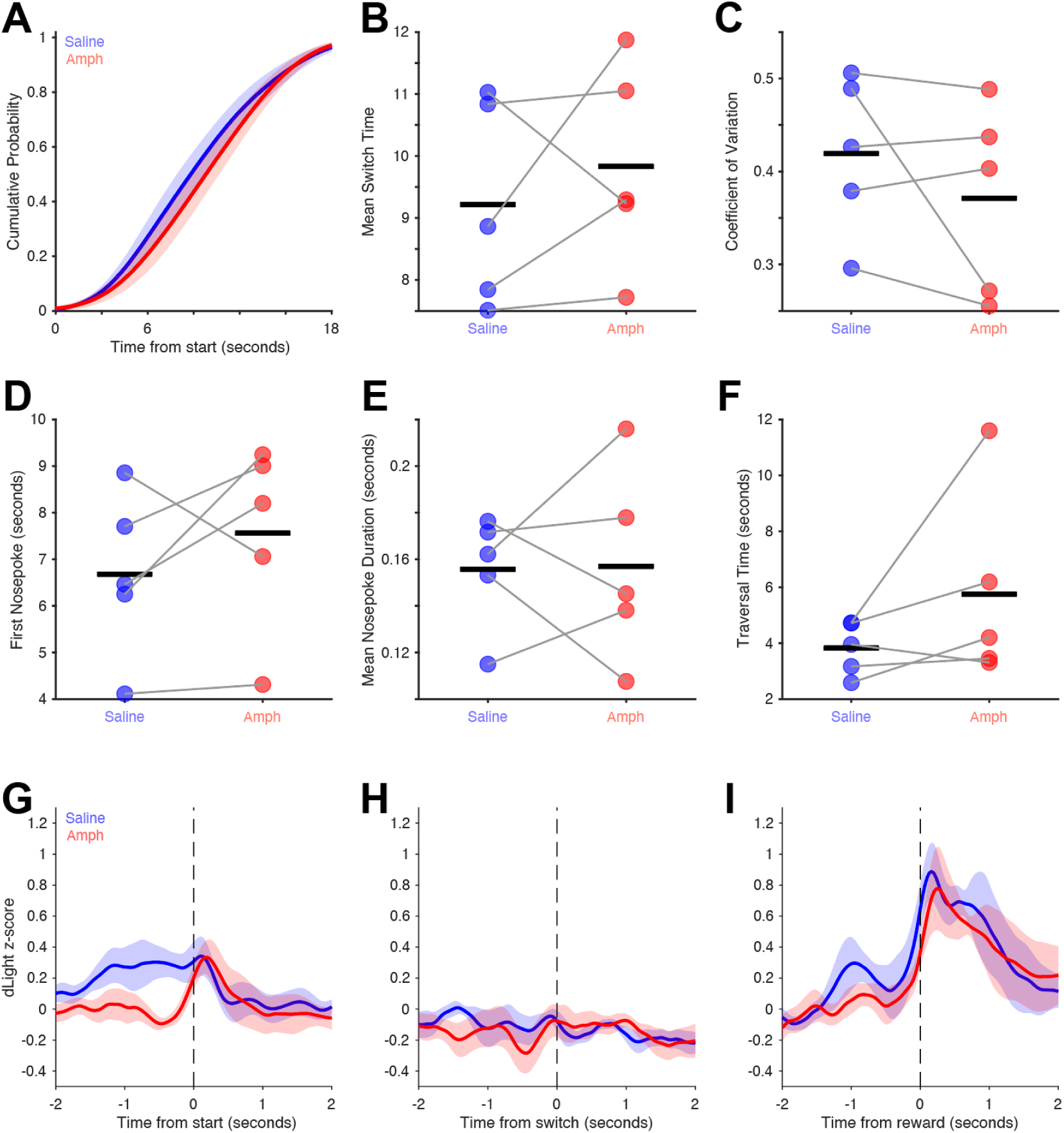
Amphetamine and dLight. (A) Cumulative probability distribution of response times from trial start for saline (blue) and amphetamine (red); shaded regions indicating the standard error of the mean (SEM). (B) Average switch time, (C) switch time coefficient of variation, (D) time of the first nosepoke, (E) average nosepoke duration, and (F) traversal time between short and long nosepokes. (G–I) Average dopamine signal measured via dLight fluorescence (z-score) aligned to G) trial start, H) switch response, and I) reward. Solid lines represent the mean z- scorewith shaded regions denoting the SEM. Data from 5 animals given saline and then amphetamine.

Finally, we were interested in how amphetamine changed dLight dynamics over the interval. We found differences when sorting dLight activity by switch times in saline vs. amphetamine sessions (Fig 6A-B). A linear mixed-effects model revealed dLight activity appeared to drop faster after the cue with amphetamine compared to saline sessions, with main effects for Time (F_(1, 4848)_ = 36.6, *p* =1X10^-9^) and a trend towards a main effect of drug (F_(1, 4848)_ = 3.4, *p* =0.06; Fig 6C). Strikingly, amphetamine interacted with the relationship between dLight slope between 0 and 9.8 seconds and switch time (main effect of slope: F_(1, 265)_ = 7.9, *p* =0.005; no effect of drug, but a significant interaction between slope and drug: F_(1, 4848)_ = 4.6, *p* =0.03). Indeed, while there was significant correlation in the saline condition between dLight slope and switch time (r=0.26, p=0.001), this was attenuated with amphetamine (r=0.01, p=0.89). Taken together, these data suggest that amphetamine changes dorsomedial striatal dLight dynamics during interval timing and provide insight into cognitive dopamine dynamics in the basal ganglia.

**Figure 6:**
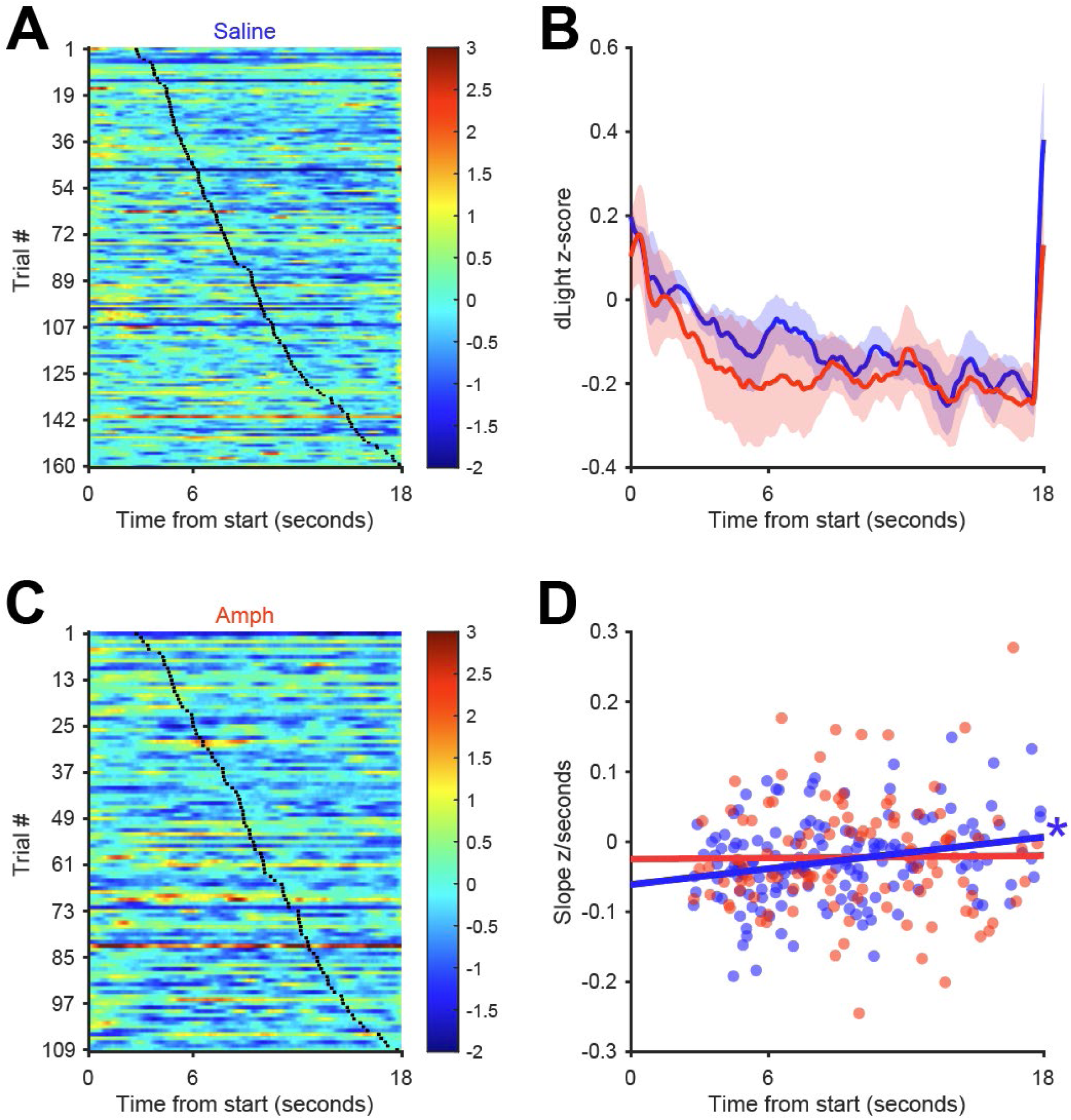
dLight dynamics in saline and amphetamine conditions. Heatmaps depicting trial-by-trial z-scored dLight fluorescence for (A) saline and (B) amphetamine sessions. Trials on the y-axis are sorted by switch time. (C) Average dLight activity for saline (blue) and amphetamine (red) conditions. Shaded regions represent standard error of the mean. (D) Scatter plot showing the relationship between the overall dLight slope between 0 and 9.8 seconds and switch time (seconds) for individual saline (blue dots) and amphetamine (red dots) trials. Solid lines represent linear regression for corresponding colors. The blue asterisk (*) denotes p < 0.05 for the saline group.

## DISCUSSION

We examined dopamine dynamics during interval timing using the fluorescent sensor dLight1.3b. We report 3 primary results. First, dLight was modulated during the interval, and ramped down from trial start to switch responses. Second, dLight slope predicted switch time, which represents animals’ internal estimates of elapsed time. Third, this relationship is distinct in males vs females, and was attenuated by amphetamine, which modulated dopamine reuptake. These data provide new information on striatal dopamine during interval timing.

Dopamine has long been known to be a highly dynamic signal, prominently modulated by reward^36^. Peaks of dopamine signals have predicted temporal estimates of reward delivery^24^. Dopamine has been described to ramp upwards representing progress towards a goal^22^, and dopamine signals have been shown to ramp up over seconds towards self-timed movements^21,23^. By contrast, we harnessed an interval timing task with fixed temporal contingencies and reward delivered after 18 seconds if animals switch from short to long nosepokes at ∼9 seconds on long correct trials. The advent of fluorescent dopamine sensors has enabled direct subsecond quantification of relative dopamine levels in target brain structures such as the dorsomedial striatum, which has prominent dopamine input^18–20,37^. While we observed prominent dopamine transients that ramped up towards reward, during the interval when mice were actively estimating time, we report evidence that dopamine ramps down, and that this downward ramp predicts animals’ internal estimates of time. Additionally, we report that males and females display distinct dopamine dynamics during interval timing where timing performance is similar, while the downward ramping dynamics across time differ by sex.

Dopamine can modulate the activity of dorsomedial striatal MSNs^38^. Past work has demonstrated that dorsomedial striatal medium spiny neurons (MSNs) expressing dopamine receptors prominently encode time^13,39,40^. Three models have been proposed, including temporal codes based on oscillatory activity, temporal basis functions, or ramping activity^41,42^. During this exact version of mouse interval timing, our work has found prominent striatal ramping activity that cannot be readily explained by movement or reward anticipation^15^. Furthermore, D1- dopamine-receptor-expressing MSNs with excitatory dopaminergic modulation ramp down, while D2-dopamine-receptor-expressing MSNs with inhibitory dopaminergic modulation ramp up. Future studies will further establish dLight dopamine dynamics during interval timing.

Our work has several limitations. First, interval timing involves working memory and attention, but does not capture other domains such as cognitive flexibility, response inhibition, or reinforcement learning. Second, fiber photometry with dLight measures relative changes in fluorescence rather than absolute dopamine concentrations. The z-scored signals reflect increases and decreases relative to session baseline and do not provide direct quantification of extracellular dopamine levels. Overall changes in dopamine concentration could be influencing dopamine dynamics across interval timing. More investigations into how overall dopamine concentration affects interval timing will be required to further understand how dorsal striatal dopamine dynamics are modulated during interval timing. Third, fiber photometry measures bulk dopamine signaling within the recorded volume and reflects a population-level signal. Finally, these findings are correlational in a single mouse strain and age range. It remains unknown whether these dopamine-behavior relationships generalize, which will be explored in future studies.

In summary, we demonstrate that DMS dopamine dynamics ramp down during interval timing. This work helps understand how striatal dopamine supports cognitive computations, which is helpful for generating new markers and targeted therapies for diseases of disrupted dopamine.

## Notes

### Competing Interest Statement

The authors have declared no competing interest.

### Funding

- National Institute of Mental Health: 5R01MH116043-05

